# Is adiposity associated with white-matter microstructural health and intelligence differently in men and women?

**DOI:** 10.1101/2022.08.20.504656

**Authors:** Arjun Patel, Jordan A. Chad, J. Jean Chen

## Abstract

The role of vascular risk in age-related brain degeneration has long been the subject of intense study. As a sub-category of vascular risk, obesity has an increasingly recognized role in influencing brain health and health-care strategies, but its association with brain health remains under-studied. Notably, no prior study has addressed sex differences in the association between adiposity and white-matter microstructural integrity, an important early marker of brain degeneration, despite known sex differences in fat storage and usage. This study focuses on the associations between adiposity (abdominal fat ratio: AFR, and liver proton density fat fraction: PDFF) and brain microstructural health (measures of white-matter microstructure using diffusion-tensor imaging, DTI). We found that fluid intelligence and reaction time are indeed associated with body fat differently in men and women. We also found significant differences in the associations of AFR with DTI metrics between sexes. These sex differences are mirrored in the associations of SBP and age with DTI metrics. Moreover, these sex differences in the AFR and SBP associations with DTI metrics persist when controlling for age. Taken together, these findings suggest that there are inherent sex-driven differences in how brain health is associated with vascular risk factors such as obesity.

## Introduction

The white matter of the human brain facilitates higher-level cognitive functioning by providing a highly connected structural network (1). White-matter microstructure exhibits unique sensitivity to changes in brain function (2), and precedes structural brain changes in disease (3, 4). White-matter microstructural degeneration in aging and disease is commonly measured using diffusion-tensor imaging (DTI) metrics, and decreases in fractional anisotropy (FA) and increases in diffusivity (MD) are commonly observed as markers of age-related microstructural degeneration (5, 6). The MD increase is typically dissected into increasing radial diffusivity (RD) and less reliable observations of changes in axial diffusivity (AD).

Prior research demonstrates links between deteriorating cognitive function and adiposity in middle-aged adults (7, 8), as well as an inverse relationship between obesity and IQ throughout the lifespan (9). Obesity is associated with excessive accumulation of adipose tissue, and has also been variably associated with reduced cortical volume, more strongly in frontal regions and less so in occipital and temporal regions (10). Furthermore, visceral obesity has recently been linked to poor white-matter microstructural health in otherwise healthy adults, as reflected by MD and FA (11) (see recent review by Okudzhava et al. (12)). Moreover, the incidence of non-alcoholic fatty-liver disease, which is closely related to obesity, has been linked with smaller total cerebral brain volumes. This finding remained statistically significant after controlling for health and lifestyle parameters, including visceral adipose tissue (13). However, the contribution of obesity to dementia in aging older adults remains unclear (7, 14, 15), partially driven by the large variation in the metrics used to quantify obesity, ranging from body-mass index (BMI) to waist-to-hip ratio (WHR). Though used in the majority of studies addressing obesity and white-matter microstructure (12), these metrics cannot be equated to level of adiposity.

Visceral adiposity is documented to account for 65-75% of the risk for human primary hypertension (manifesting commonly as elevated BP) (16, 17). Related vascular risk factors such as arterial stiffness (18–20), high blood glucose (21), diabetes mellitus (22, 23), hypercholesterolaemia (Haley et al., 2018), and anthropometric indices (e.g. body mass index; BMI) (21) are also known to modify the normal aging process of the white-matter microstructure (24). Exposure to these vascular risk factors may exacerbate these white-matter microstructural changes (25),preferentially in anterior compared to posterior brain regions (24). It is well established that high abdominal fat (adiposity) and blood pressure (BP) are interrelated vascular-risk factors (26). They both contribute to pathological aging, such as in the case of type 2 diabetes (27). Unlike diastolic BP (DBP), systolic BP (SBP) increases with age, and is more predictive of stroke and heart disease (28). Elevated SBP and cholesterol are associated with cognitive decline and neurodegeneration (29, 30). Previous literature has found associations of both adiposity and SBP with human white matter microstructure by using metrics derived from DTI (31). SBP and adiposity have been shown to be negatively correlated with FA and AD and positively correlated with MD and RD in regions including the corpus callosum (31–37), fronto-occipital fasciculi (31, 33, 38), splenium (36), fornix (36, 39), corona radiata (37), cingulum (38), and cerebellar peduncles (37).

A growing number of studies have demonstrated that the effects of aging, adiposity and high SBP on brain health can vary between sexes. Different vascular risk factors, such as hypertension, heart disease and high homocysteine in men, and smoking status in women (40), were found to be associated with occurrence of white matter small-vessel disease in both sexes. One study of BMI found significantly lower AD among males and females with higher BMI in the corpus callosum, and significantly higher RD among only females with higher BMI (34), which suggests a sex-dependence. In a study on brain volume changes, Armstrong et al. (41) demonstrated that hypertension and higher HDL cholesterol were protective against volume loss in males, whereas hypertension was associated with steeper decline in gray matter for females and obesity was protective against volume loss in temporal gray matter. Non-cognitive studies such as brain atrophy studies noted differences in symmetry of atrophy between men and women with age, suggesting differences in susceptibility (42). SBP has been found to increase more quickly with age in men than women longitudinally (43). Similarly, arterial stiffness has been found to be more strongly associated with executive function in men than in women (44). Contradictorily, however, pulse-wave velocity - a close correlate of hypertension and arterial stiffness - has been shown to increase more with age in women than men (45, 46). Moreover, age-related cerebral blood flow (CBF) increases in the cerebellum were significantly higher in women, while age-related CBF increases in the occipital lobe were greater in men (47). Nonetheless, existing studies on the sex-dependence of these associations are limited in sample size and methodological rigor.

Given the prevalence of obesity in contributing to vascular risk, it is natural to investigate whether the unique interactions between obesity measures and brain microstructure underlies these reported sex differences, especially as body fat content strongly depends on sex. Nonetheless, the majority of existing studies cited earlier infer adiposity from BMI and WHR, which can differ in interpretation between sexes, and are poor surrogates of obesity. Previous studies have also theorized a link between the expression of female hormones such as estrogen and a higher predisposition towards abdominal obesity. Women are known to have higher percentage body fat than men, and store proportionately more fat in the gluteal-femoral rather than the abdominal (visceral) region (48). Women also metabolize fat differently, reflected by reports of lower resting fat oxidation in both younger and older women (49, 50). This sex differences in hepatic fat oxidation and synthesis may underlie the higher incidence of fatty liver disease in men (51), which is strongly associated with obesity (52, 53). However, when experiencing estrogen decline in later years, women are more predisposed to metabolic diseases such as type 2 diabetes (39), which has a well-known negative impact on white-matter microstructural health (25, 54).

In light of these theorized sex differences and the current lack of rigorous investigations into sex differences in the association between obesity and brain microstructure, the present study seeks to fill this knowledge gap by using the comprehensive measures of obesity available from the Uk Biobank. We also investigate whether vascular-risk influences on white matter are distinct from those of aging, and whether this influence is more pronounced in women than in men. We further relate the effects of adiposity to those of other vascular risk factors to demonstrate their similarities and differences in terms of sex differences.

## Methods

### Study Participants and Anthropometrics

All participants were part of the UK Biobank initiative (ukbiobank.co.uk), and all data were obtained as a part of the UK Biobank Application 40922. Subjects were excluded from our study if they reported a neurological, psychological or psychiatric disorder, neurological injury, or history of stroke. We selected an initial sample of 700 subjects to cover the entire UK Biobank age range at regular intervals, with equal number of males and females in each decade. Our non-imaging analysis involves metrics of body composition and of vascular risk. We also included metrics of intelligence, specifically in terms of reasoning.

Body composition metrics:

⍰ Abdominal fat ratio (AFR): total abdominal fat divided by total abdominal fat and thigh muscle volume, assessing the distribution between fat and muscle volume;
⍰ Abdominal subcutaneous adipose tissue volume normalized by height (ASAT): a measure of obesity that has been related to varying degrees of cardiometabolic risk (55);
⍰ Body mass index (BMI): defined as the body weight in kg divided by height squared (m^2^), BMI has been widely used as a surrogate for obesity (9, 56);
⍰ Liver proton-density fat fraction (PDFF): a measure related to fatty liver disease that is weakly associated with ASAT (57);
⍰ Total abdominal adipose tissue index (TAATi): the total abdominal fat volume normalized by height squared (L/m^2^), a fat-specific version of BMI that is associated with the incidence of metabolic syndromes (58).

Metrics of vascular risk:

⍰ Systolic blood pressure (SBP): the association between elevated SBP and cognitive decline has been inconsistent; a recent study established this link in older women (59);
⍰ Diastolic blood pressure (DBP): elevated DBP, alongside SBP, is associated with poorer cognitive performance in middle-to-older men (60).
⍰ Arterial pulse-wave stiffness index (PWV): This is measured as the time between peaks of the pulse wave divided into the person’s height. Arterial stiffness index is a well-established correlate of cerebrovascular health and cognition (20, 61, 62), and has been closely associated with SBP (63–65).

Metrics of intelligence (66):

⍰ Fluid intelligence (FI): the capacity to solve problems that require logic and reasoning ability, independent of acquired knowledge;
⍰ Reaction time (RT): a measure of processing time in a cognitive task (e.g. a card game).

After ensuring adequate quality of diffusion MRI data from all participants, we performed quality assessment of the data to include only participants with complete sets of anthropometrics. The resultant sample consisted of 281 subjects aged 46-79 (158 male, 128 female).

### Diffusion MRI Data

Diffusion MRI data were acquired as part of the UK Biobank on a Siemens Skyra 3T system with 5 b=0 and 50 b=1000s/mm^2^ at TR = 3.6s, TE=92ms, matrix size=104×104×72 with 2×2×2mm^3^ resolution, 6/8 partial Fourier, 3x multislice acceleration and no in-plane acceleration. An additional 3 b=0 volumes were acquired with a reversed phase encoding. The data were corrected for eddy-current and susceptibility-related distortions via EDDY^5^ and TOPUP^6^, respectively.

### Anthropometric Statistical Analysis

The associations across age, body-composition metrics, vascular risk metrics and intelligence are assessed using Spearman’s correlation. Metrics of body composition that are significantly associated with intelligence are identified for further analysis. Likewise, for comparison, metrics of vascular risk that are significantly associated with intelligence are also identified from this analysis.

### Imaging Statistical Analysis

FSL’s Tract-Based Spatial Statistics (TBSS) was used to obtain a WM skeleton.^8^ For voxelwise analysis, an effect was computed using a linear model defined by p<0.05 with correction for multiple comparisons (cluster-wise controlling family-wise error rate with 500 permutations) and controlling for age as per FSL’s randomise. ^9^

The following DTI metrics were used in analyses: FA, MD, AD and RD. Each DTI metric was investigated for positive and negative associations with either SBP or AFR. This was done without controlling for sex or age, controlling for only sex, and controlling for both sex and age. Age was controlled to account for age-related changes in white-matter microstructure.

In addition, a comparison of the associations between either indicators of obesity and the DTI metrics was conducted for males and females. This was conducted without controlling for age, and while controlling for age to account for age-related changes in white-matter microstructure. The same was done with indicators of vascular risk for reference.

## Results

**Table 1.** Participant demographics, body composition, vascular and cognitive measures. AFR=abdominal fat ratio; ASAT=abdominal subcutaneous adipose tissue volume (L/m^2^); BMI = body-mass index; DBP=diastolic blood pressure (mmHg); FI=fluid intelligence; PDFF=liver proton density fat fraction; PWV=pulse-wave stiffness index; RT=reaction time (msec); SBP=systolic blood pressure (mmHg); TAATi=total abdominal adipose tissue index (L/m^2^).

| Metric | Women | Men | Combined |
| --- | --- | --- | --- |
| Age (years) | 60.4±9.7 | 61.4±9.6 | 60.9±9.6 |
| AFR | 0.52±0.10 | 0.44±0.10 | 0.48±0.11 |
| ASAT ($L/m^2$) | 7.6±3.2 | 6.2±3.0 | 6.8±3.2 |
| TAATi ($L/m^2$) | 3.8±1.6 | 3.6±1.6 | 3.7±1.6 |
| BMI | 25.4±4.0 | 27.5±4.7 | 26.5±4.5 |
| PDFF | 3.1±3.4 | 5.3±5.3 | 4.3±4.7 |
| SBP (mmHg) | 132.8±19.1 | 141.3±16.6 | 137.4±18.3 |
| DBP (mmHg) | 76.5±10.4 | 88.3±11.1 | 78.6±10.9 |
| PWV (cm/s) | 8.4±2.7 | 9.8±2.3 | 9.1±2.6 |
| FI | 6.7±1.9 | 6.9±2.0 | 6.8±1.9 |
| RT (ms) | 603.7±119.0 | 565.7±119.4 | 583.1±120.5 |

The BMI threshold for being overweight is 25, and for obesity is 30 (67). Of the 128 female subjects, 61 had a BMI of greater than 25 (47.7%), and 15 (11.7%) had a BMI greater than 30. Of the 153 male subjects, 98 (64.1%) had a BMI of greater than 25, and 32 (20.9%) had a BMI greater than 30.

The cross-correlation across age, body-composition, vascular-risk and intelligence-related variables are shown for men and women separately in **Fig. 1** and **2**. The significant correlations are summarized in **Fig. 3**. Since it is unclear whether the correlations should be assumed linear, we used Spearman’s rank correlation. Nonetheless, all significant correlations appear linear. As shown in **Fig. 3**, Note that only AFR is associated with both FI and RT, but differently between males and females. That is, AFR is associated with FI in men only, and with RT in women only. RT in females (not in males) is additionally associated with SBP and AMRA.

**Figure 1.**
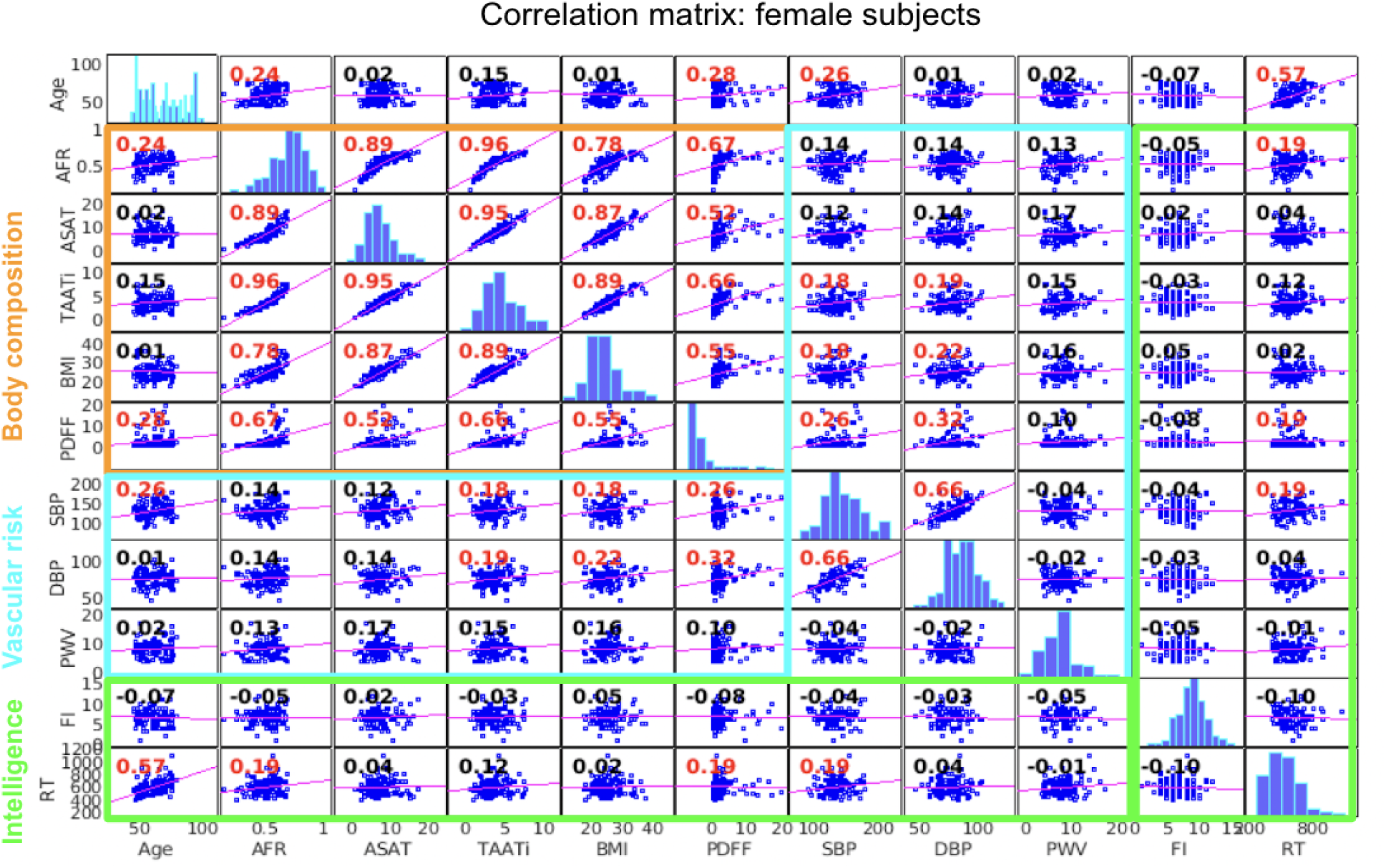
Data correlation matrix for female subjects. The Spearman correlation plots are shown for each pair of variables, with the correlation coefficients are indicated in each plot, and significant correlations indicated in red. The histograms of parameters across subjects are shown on the diagonal line. The strongest associations are: (i) AFR, TAATi, ASAT, BMI and PDFF; (ii) age and RT. Age=in years; AFR=abdominal fat ratio; ASAT=abdominal subcutaneous adipose tissue volume (L/m^2^); BMI = body-mass index; DBP=diastolic blood pressure (mmHg); FI=fluid intelligence; PDFF=liver proton density fat fraction; PWV=pulse-wave stiffness index; RT=reaction time (msec); SBP=systolic blood pressure (mmHg); TAATi=total abdominal adipose tissue index (L/m^2^).

**Figure 2.**
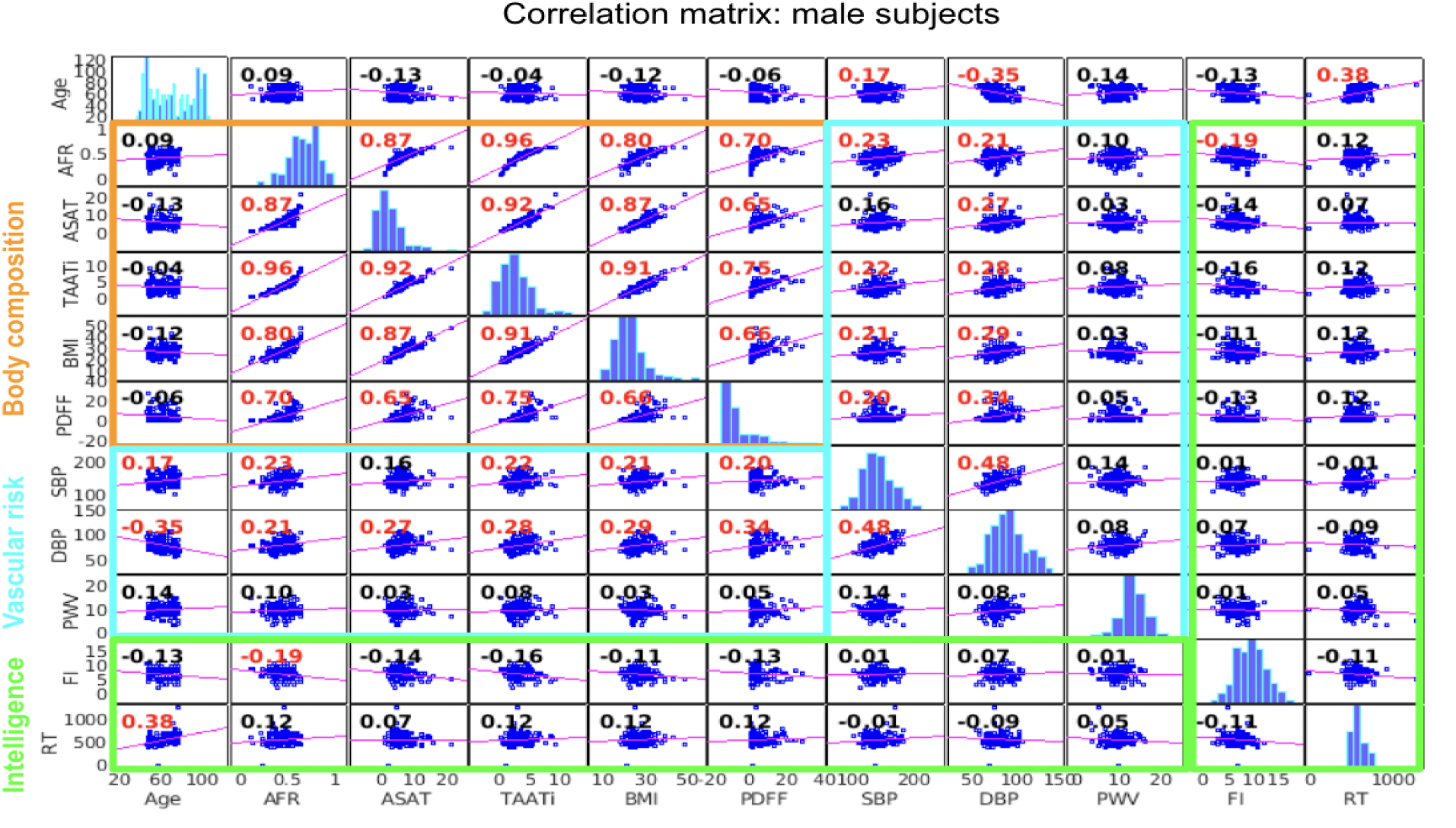
Data correlation matrix for male subjects. The Spearman correlation plots are shown for each pair of variables, with the correlation coefficients are indicated in each plot, and significant correlations indicated in red. The histograms of parameters across subjects are shown on the diagonal line. The strongest associations are: (i) AFR, TAATi, ASAT, BMI and PDFF; (ii) age and RT. Age=in years; AFR=abdominal fat ratio; ASAT=abdominal subcutaneous adipose tissue volume (L/mm^2^); BMI = body-mass index; DBP=diastolic blood pressure (mmHg); FI=fluid intelligence; PDFF=liver proton density fat fraction; PWV=pulse-wave stiffness index; RT=reaction time (msec); SBP=systolic blood pressure (mmHg); TAATi=total abdominal adipose tissue index (L/m^2^).

**Figure 3.**
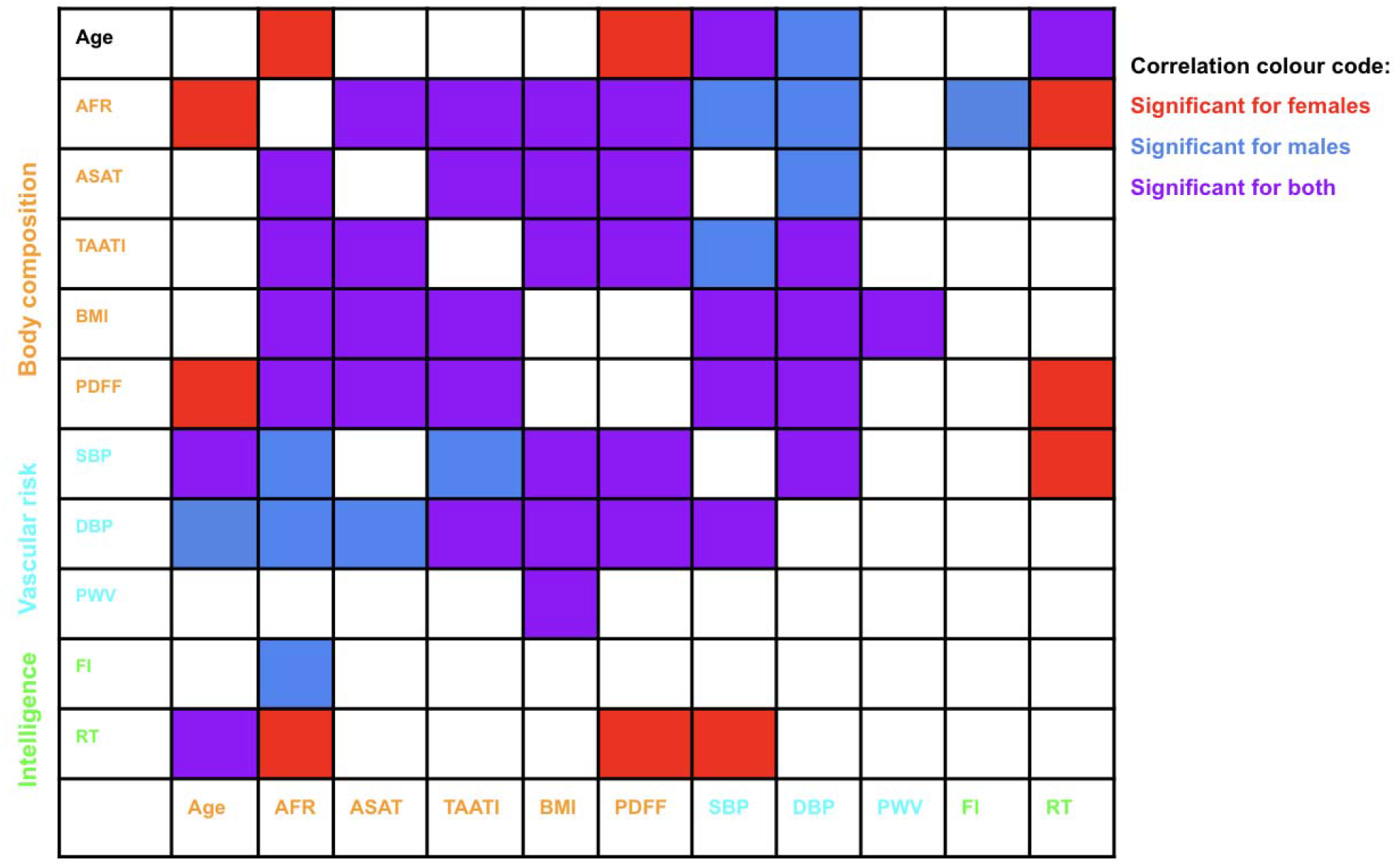
Data correlation matrix comparisons between female and male subjects. Intelligence-related variables are more strongly associated with obesity and vascular-risk factors in females than in males, while intelligence in both groups are associated with age. Age=in years; SBP=systolic blood pressure (mmHg), DBP=diastolic blood pressure (mmHg); AFR=abdominal fat ratio; PWV=pulse-wave stiffness index; ASAT=abdominal subcutaneous adipose tissue volume; TAATI=total abdominal adipose tissue index; PDFF=liver proton density fat fraction; FI=fluid intelligence; RT=reaction time (msec).

### Body composition, vascular risk and age

AFR and SBP were significantly and positively correlated with age in females only (p<0.05) (**Fig. 1a** and **1b**, respectively). No significant correlation between AFR or SBP and age was observed amongst males.

### Associations between body fat and DTI parameters

We first assessed the association between DTI parameters and age, as well as sex differences in these age associations. The results are reported in **Fig. A1-2** in Supplementary Materials, respectively, and the age associations in particular are similar to those reported previously using a similar UK Biobank dataset (68, 69).

As shown in **Fig. 5a**, AD alone is negatively associated with AFR, mainly in the genu of corpus callosum, forceps minor, pontine crossing tract (a part of MCP), corticospinal tract, cerebral peduncle, right anterior thalamic radiation, and right superior corona radiata. After controlling for sex differences (**Fig. 5b**), both AD and FA are negatively associated with AFR, but in different locations; the AD-AFR associations are found in the pontine crossing tract, corticospinal tract, and cerebral peduncle, while the FA-AFR associations are found in the body and genu of the corpus callosum, forceps major and minor, fornix, anterior thalamic radiation, superior longitudinal fasciculus, left corticospinal tract, left external capsule, left cingulum, left inferior longitudinal fasciculus, left inferior fronto-occipital fasciculus, left anterior corona radiata, left posterior corona radiata, and left superior corona radiata. Once age and sex are both controlled for, FA is no longer associated with AFR (**Fig. 5c**). No significant positive associations are observed in any of the metrics, and no significant associations of either kind is observed in RD or MD. **Fig. A3** in Supplementary Materials corresponds to scatterplots corresponding to the region of interest (ROI) in which FA is significantly associated with AFR, controlling for sex (**Fig. 5b**).

**Figure 4.**
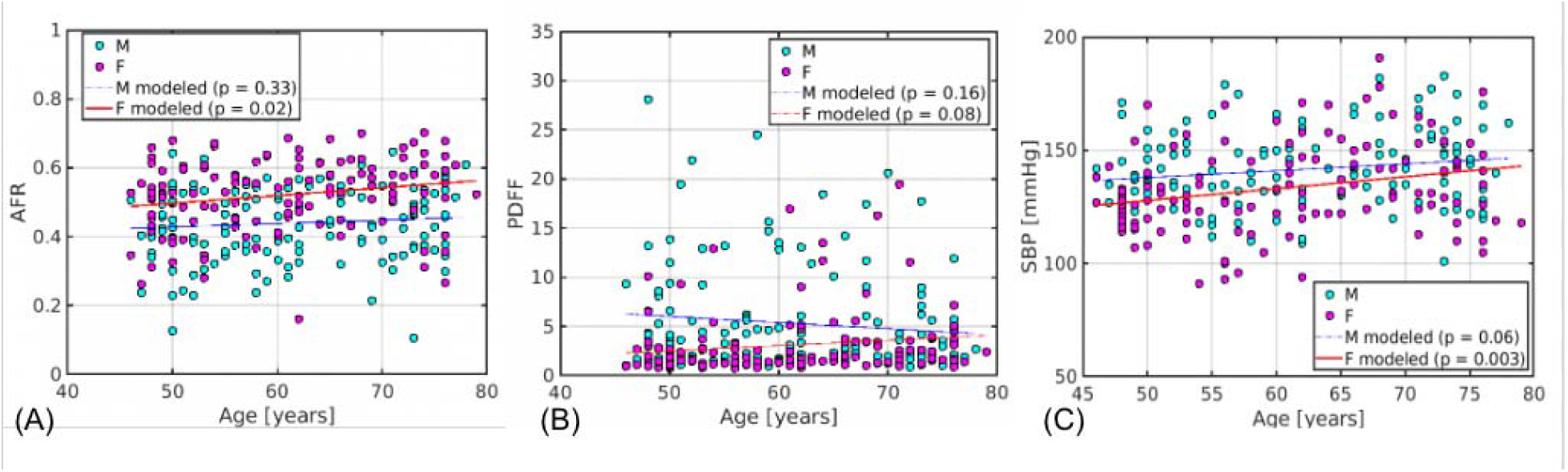
AFR, PDFFSBP plotted against age for males and females. (a) AFR was only significantly associated with age in females. (b) PDFF is not significantly associated with age in either males or females, and is associated with remarkably large variability in men. (c) SBP was only significantly associated with age in females. The thinner lines represent model fits that failed to achieve significance at the 0.05 level.

**Figure 5.**
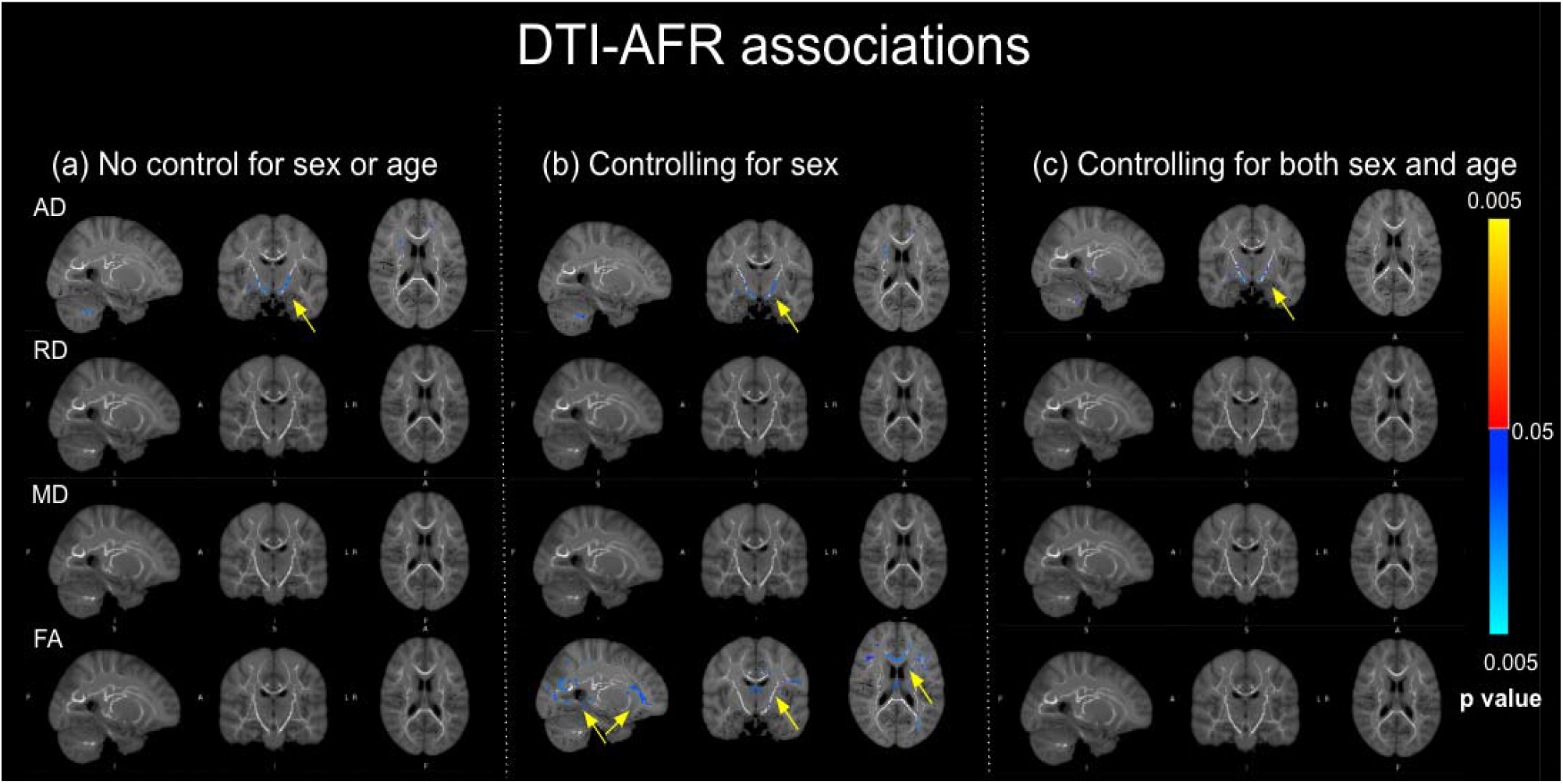
DTI-AFR associations. Blue indicates significant negative associations (further highlighted by yellow arrows for better visibility), corrected for multiple comparisons. The corrected p value of the association is shown in the colourbar. There are no positive associations observed in any of the parameters.

No significant association was found between PDFF and DTI parameters, with or without controlling for age and sex.

### Associations between blood pressure and DTI parameters

AD, RD and MD were all positively associated with SBP (**Fig. 6a**), although their areas of associations differed. AD was positively associated with SBP in the posterior corpus callosum, right anterior and posterior limbs of the internal capsule, external capsule, anterior thalamic radiation, uncinate fasciculus, inferior fronto-occipital fasciculus, forceps major and minor, corticospinal tract, superior longitudinal fasciculus and throughout the corona radiata. In contrast, RD-SBP associations were more anterior, and MD-SBP associations are the most extensive of all three, found in the body and genu of the corpus callosum, anterior corona radiata, posterior corona radiata, superior corona radiata, corticospinal tract, left external capsule, superior longitudinal fasciculus, posterior limb of internal capsule, anterior thalamic radiation, and superior fronto-occipital fasciculus. All of these associations were weakened after controlling for sex (**Fig. 6b**), and further weakened after controlling for both sex and age. There was no negative association with SBP in any of the metrics, and FA was not significantly associated with SBP in any condition. **Fig. A4** in Supplementary Materials corresponds to scatterplots corresponding to the region of interest (ROI) in which MD is significantly associated with AFR, controlling for sex (**Fig. 6b**).

**Figure 6.**
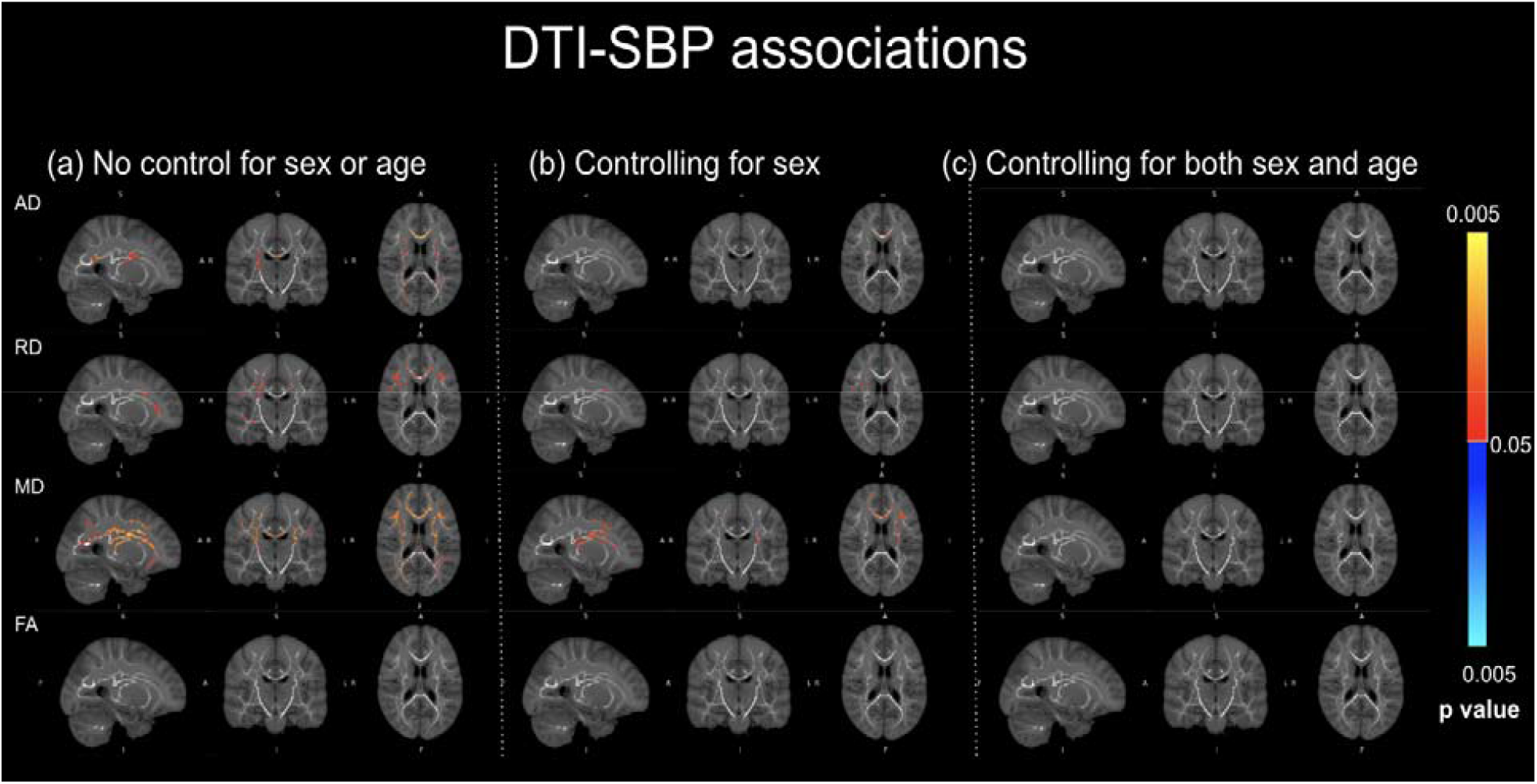
DTI-SBP associations. Orange indicates significant positive associations, corrected for multiple comparisons. The corrected p value of the association is shown in the colourbar. There are no negative associations observed in any of the parameters.

### Sex differences in DTI-body-fat associations

The magnitude of the associations of AFR with RD and FA were significantly greater in females than in males (**Fig. 7a**). In the case of AD, the differences are found in the genu, body and splenium of corpus callosum, forceps minor, superior corona radiata, posterior corona radiata, superior longitudinal fasciculus, posterior limb of internal capsule, external capsule, inferior fronto-occipital fasciculus, inferior longitudinal fasciculus, and left cerebral peduncle. In the case of RD, the differences are found in the body, genu, and splenium of corpus callosum, forceps minor, superior corona radiata, posterior corona radiata, superior longitudinal fasciculus, posterior limb of internal capsule, external capsule, inferior fronto-occipital fasciculus, inferior longitudinal fasciculus, and left cerebral peduncle. After controlling for age, the female group still exhibited stronger DTI-AFR associations, but the sex difference is noticeably diminished (**Fig. 7b**). No sex difference in AFR associations were found for AD and MD.

**Figure 7.**
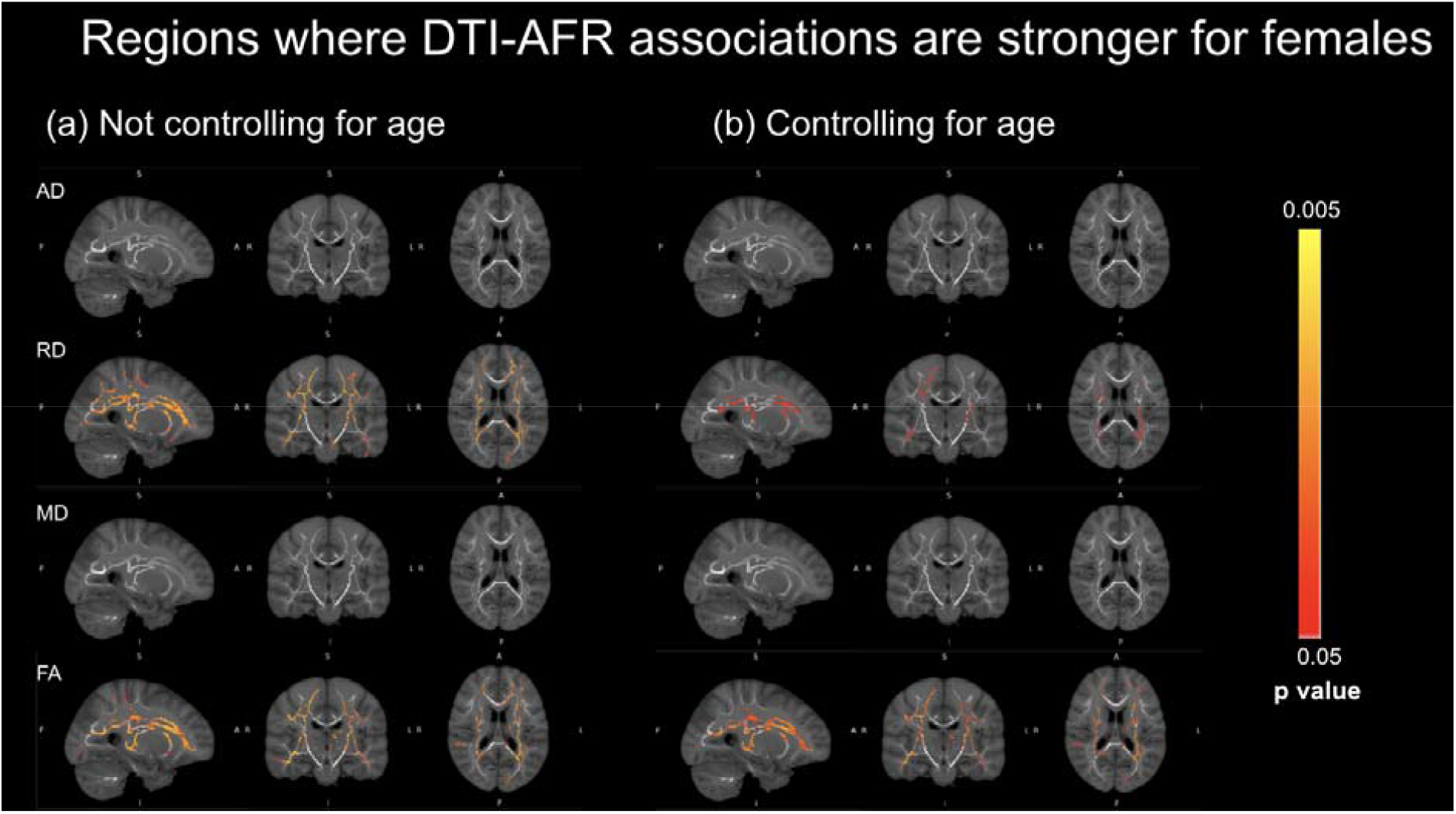
Regions where DTI-AFR associations are stronger for females than males. Displayed are differences in absolute associations. Orange indicates significantly stronger absolute associations in female subjects, corrected for multiple comparisons. The corrected p value of the association is shown in the colourbar. There are no regions in which the male subjects exhibited a stronger association.

Though no significant associations between PDFF and DTI parameters were found, PDFF is more strongly associated with DTI parameters of white-matter microstructural integrity in women (**Fig. 8**). Again, displayed are differences in absolute associations. The sex differences are removed by controlling for age.

**Figure 8.**
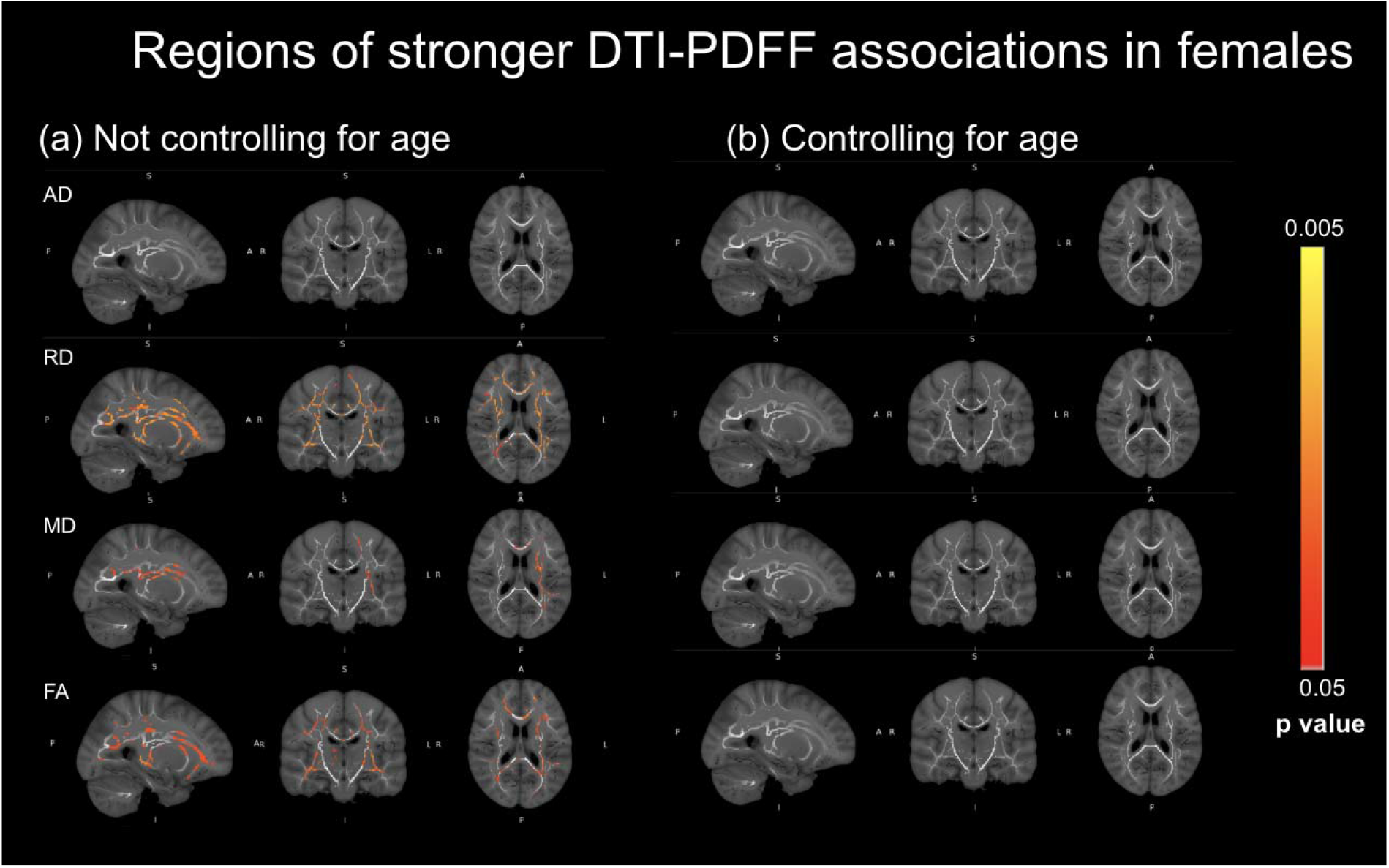
Regions where DTI-PDFF associations are stronger for females than males. Displayed are differences in absolute associations. Orange indicates significantly greater effects in female subjects, corrected for multiple comparisons. The corrected p value of the association is shown in the colourbar. There are no regions in which the male subjects exhibited a stronger association.

**Figure 9.**
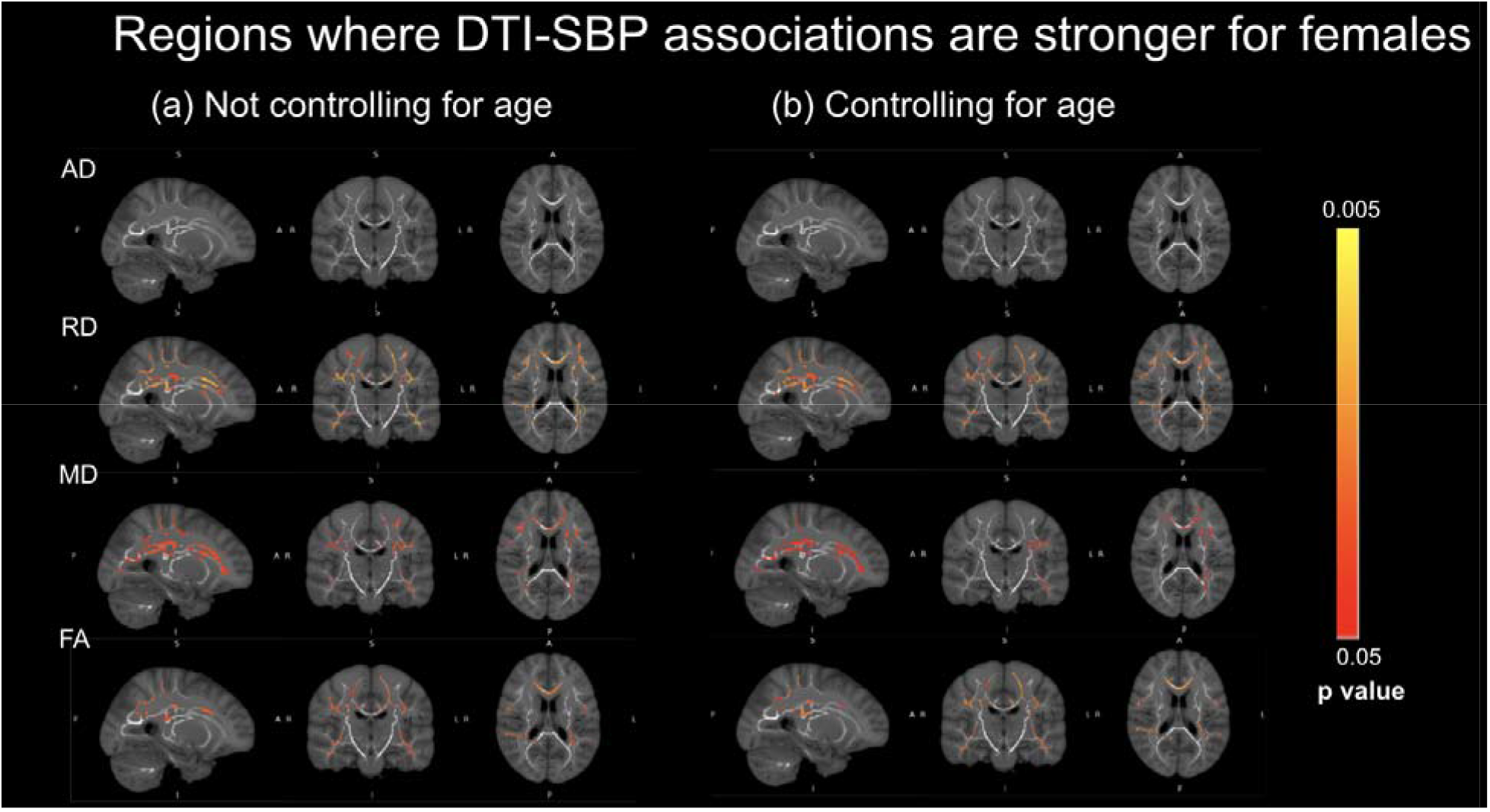
Regions where DTI-SBP associations are stronger for females than males. Orange indicates significantly greater effects in female subjects, corrected for multiple comparisons. The corrected p value of the association is shown in the colourbar. There are no regions in which the male subjects exhibited a stronger association.

### Sex differences in DTI-SBP associations

Likewise, the magnitude of the associations of SBP with RD, FA, and MD were significantly greater in females than in males (**Fig. 5a**). For all three parameters, the sex difference is found in the body, genu, and splenium of corpus callosum, corticospinal tract, superior longitudinal fasciculus, superior corona radiata, anterior corona radiata, posterior corona radiata, inferior fronto-occipital fasciculus, inferior longitudinal fasciculus, forceps minor, forceps major, and anterior thalamic radiation. These differences were only negligibly altered by controlling for age (**Fig. 5b**). No sex difference in absolute associations were found for SBP and AD.

## Discussion

In the past decade, the role of vascular risk factors in age-related brain degeneration has long been the subject of intense study (70–72). As its own sub-category of vascular risk, adiposity has an increasingly recognized role in influencing brain health and health-care strategies, but its association with brain health remains under-studied. Notably, no prior study has addressed the association between metrics of obesity and white-matter microstructural integrity, an important early marker of brain degeneration (73–75). Given known sex differences in fat storage and usage, this study focuses on the associations between markers of body fat (abdominal fat ratio: AFR, and liver proton density fat fraction: PDFF) and brain microstructural health, shedding light also on the association between body fat and intelligence metrics. To help us position these findings in terms of sex differences in general vascular risk, we also included systolic blood pressure (SBP) as a reference vascular-risk factor, and age as a reference variable. We found significant differences in the associations of AFR with DTI metrics between sexes. These sex differences are also mirrored in the association between SBP and DTI metrics. Moreover, these sex differences exist irrespective of age. Taken together, these findings suggest that there are inherent sex-driven differences in how brain health is associates with vascular risk factors such as obesity.

### Literature support for sex differences in vascular risk

Previous literature supports the existence of associations between vascular risk factors and white matter health as well as cognition. In particular, arterial stiffness is a known risk factor that co-interacts with hypertension (Safar et al., 2018). For instance, arterial stiffness is found to be associated with axonal integrity in the corpus callosum, the internal capsule and the corona radiata (Badji et al., 2019). Sex differences in the effect of individual risk factor has been documented in the literature, prompting our hypothesis that the respective effect of the risk factors on white-matter microstructure and cognition may be influenced by sex as well. These differences are hypothesized to be a result of sex hormone differences between males and females. For example, arterial stiffness is known to increase faster with age in females as estrogen levels drop post-menopause (DuPont et al., 2019; Mitchell et al., 2014). Recently, sex was found to moderate the associations between pulse-wave velocity (a measure of arterial stiffness; PWV) and cerebrovascular reactivity (CVR), as well as pulse-wave velocity and executive function; the effect of PWV on CVR was positive in males and negative in females, while the effect of PWV on executive function was negative in males but not significant in females (Sabra et al., 2021). Based on this evidence, age related increases in vascular risk factors themselves may be sexually dimorphic. In fact, a separate study found that there were stronger correlations between pulse pressure and restricted isotropic diffusion in late middle to early older women compared to men in that age group (Reas et al., 2021). Hence, there is strong literature support for the hypothesis that sex differences may influence the relations between vascular risk factors, white matter health and intelligence.

Evidence also supports the existence of sex differences in the association between white-matter health and secondary risk factors such as Type II diabetes (25). The length of time before diagnosis with a vascular risk factor may be associated with greater risk of WM microstructural changes in women compared to men. Furthermore, risk factors such as diabetes, mid-life obesity and hypertension have been described to increase vascular contributions to cognitive impairment and dementia (VCID) more so in females (76). In contrast, stroke, hyperlipidemia, and heart disease contribute more in males (76). Another known risk factor for white-matter degeneration is APOE status, and research found that vascular risk factors and APOE ε4 allele had interactive effects on white-matter microstructure, and that having multiple (≥2) vascular factors was particularly detrimental to white matter integrity among APOE ε4 carriers (Wang et al., 2015). The same study also associated changes in white-matter microstructure with changes in Mini-Mental State Examination decline. A study on patients with amnestic mild-cognitive impairment also found that APOE-related risk in women may be associated with decreased activity in both gray matter and white matter compared with men (Lin et al., 2020).

### Adiposity and white-matter integrity

A primary finding of this study is a significant association between body fat and DTI-based white-matter microstructural integrity. This is consistent with accumulating recent reports of visceral obesity being linked to poor white-matter health in otherwise healthy adults (11, 12, 77, 78). The affected tracts, including the genu of corpus callosum, forceps minor, pontine crossing tract (a part of MCP), corticospinal tract, cerebral peduncle, right anterior thalamic radiation, and right superior corona radiata, are associated with reward processing (79), inhibitory control (80), sensory integration (81) and memory (81, 82). Nonetheless, there remains substantial variability in the existing literature, in part due to BMI and WHR being used as surrogates of adiposity in the majority of cases (12). While these prior studies found FA to be significantly negatively associated with metrics of obesity, the AFR-FA association is just below the threshold of significance in this study, the same negative relationship can be observed (see **Fig. 3A** in Supplementary Materials). The DTI associations with AFR is stronger than those with PDFF. It is established that an early consequence of obesity is chronic low-grade inflammation, which has in turn been associated with fluid intelligence (83). Systemic inflammatory responses are associated with increased diffusivity and decreased fractional anisotropy (84, 85).

Despite a wealth of literature on sex differences in vascular risk factors, there is little research into how adiposity confers risk to brain health differently in men and women. The current study found that the sex differences in the associations of AFR with the DTI metrics, implicating a broad set of tracts including the genu, body and splenium of corpus callosum, forceps minor, superior corona radiata, posterior corona radiata, superior longitudinal fasciculus, posterior limb of internal capsule, external capsule, inferior fronto-occipital fasciculus, inferior longitudinal fasciculus, and left cerebral peduncle. These sex differences persisted even when controlling for age. Thus, even when not considering advanced aging or morbid obesity, adiposity has a more detrimental effect on the white-matter health of women, despite there being a lower fraction of obese women than obese men in this study (11.7% vs. 20.9%). Moreover, controlling for sex further revealed anterior white-matter regions in which integrity is associated with AFR. This further supports studies of the sex differences in the body-fat-brain connection. Recent animal experiments suggest that females are more resistant to obesity than males (86), but also suggest that to arrive at the same level of adiposity as in men, women may have been exposed to more unhealthy lifestyles, thus leading to increased risk for brain-tissue damage.

### Comparison with the effects of SBP

This study found that higher SBP is associated with poor white-matter microstructural integrity, which is well in accordance with prior findings (32). Moreover, sex differences in the associations of SBP with the DTI metrics persisted when controlling for age, suggesting that such differences exist even across individuals of the same age group.

An interesting difference between the SBP and AFR associations was found: SBP is associated with microstructural integrity primarily in the anterior-superior white matter. Conversely, AFR is associated with white-matter integrity primarily through AD decrease, and in the posterior-inferior white matter and this effect even survives covarying for age. This posterior-inferior region is known to exhibit AD associations with age opposite those of the anterior-superior white matter (68, 69, 87). This negative AD-AFR association can be attributed to increasing axonal debris accumulation with increasing degree of obesity, and is likely an early indicator of fibre degeneration, especially as there is an absence of mirroring effects in MD. The stronger SBP-DTI association in women do not necessarily subscribe to the same mechanisms as that of the AFR-DTI association. The stronger SBP influence in women is consistent with recent findings that women exhibit greater decline in cardiovascular health for a given level of low-grade inflammation (88).

### The role of age

In accordance with previous reports by ourselves and others (68, 69, 74, 89), we found robust age-associations in all DTI metrics (Supplementary Materials). Furthermore, we found that while controlling for age greatly reduced all associations with white-matter microstructural integrity, suggesting that body fat, like SBP, increasingly influences brain health as adults age. It is possible that aging enhances these sex differences in associations between SBP/AFR and DTI metrics of white matter health. An alternative explanation is that aging has different effects on males and females. Based on these differences, future studies investigating brain vulnerability to vascular risk factors and age should account for sex differences for their analyses.

Conversely, a previous study with participants aged between 44.23–79.41 years (meanl1=l161.97 years) observed that associations between general vascular risk and global measures of white matter health (global MD and global FA) are independent of age, pointing towards a more stable association between the two variables (Cox et al., 2019). If the strength of the association between vascular risk and white matter health does not change significantly with age, it is possible that baseline sex differences the associations between vascular risk factors and DTI metrics remain stable with age. Hence, further investigation could focus on how these sex differences change as a function of age. In addition, future investigation may explore whether sex differences are unique to each vascular risk factor, or whether they apply to additive effects of multiple vascular risk factors on WM health as well.

### Obesity and intelligence

Obesity has been linked with a decline in the intelligence quotient even within young adults (90). In our sample, aside from SBP, AFR and PDFF are the only parameters associated with intelligence (in terms of fluid intelligence (FI) and reaction time (RT)) (**Fig. 1-3**), and do so differently in the male and female groups. Taken with the findings of white-matter health, we propose that white-matter microstructural integrity, which is adversely affected by low-grade inflammation brought about by obesity, may be a key factor linking obesity and intelligence. Existing literature links body fat and hypertension to cognitive deficits such as those affecting memory and executive function (Elias et al., 2003). Lower white matter microstructural integrity in regions such as the anterior limb of the internal capsule, fornix, superior longitudinal fasciculus and superior corona radiata have previously been associated with reduced executive function (Heidi et al., 2013). In addition, changes in diffusivity within white matter lesions and normally-appearing white matter have been associated with overall worse information processing speed, global cognition and motor speed (Vernooij et al., 2009). Therefore, future studies should also explore whether sex differences in associations between vascular risk factors, white matter health, and measures of intelligence align with the sex differences observed in this study.

### Limitations

As the focus of this work is intelligence, we have not included memory assessments that are available through the UK Biobank. Indeed, in the context of age, memory is extremely relevant in a study on obesity, and will be the subject of a follow-up study. Moreover, as this is a cross-sectional study, we are unable to conclude regarding the causality of these sex-obesity interactions in influence white-matter integrity and intelligence (9, 56).

## Acknowledgments

This work has been supported by the Canadian Institutes of Health Research and the Sandra Rotman Foundation.

## Notes

### Competing Interest Statement

The authors have declared no competing interest.

