## Supplementary Materials for "Is adiposity associated with white-matter microstructural health and intelligence differently in men and women?"

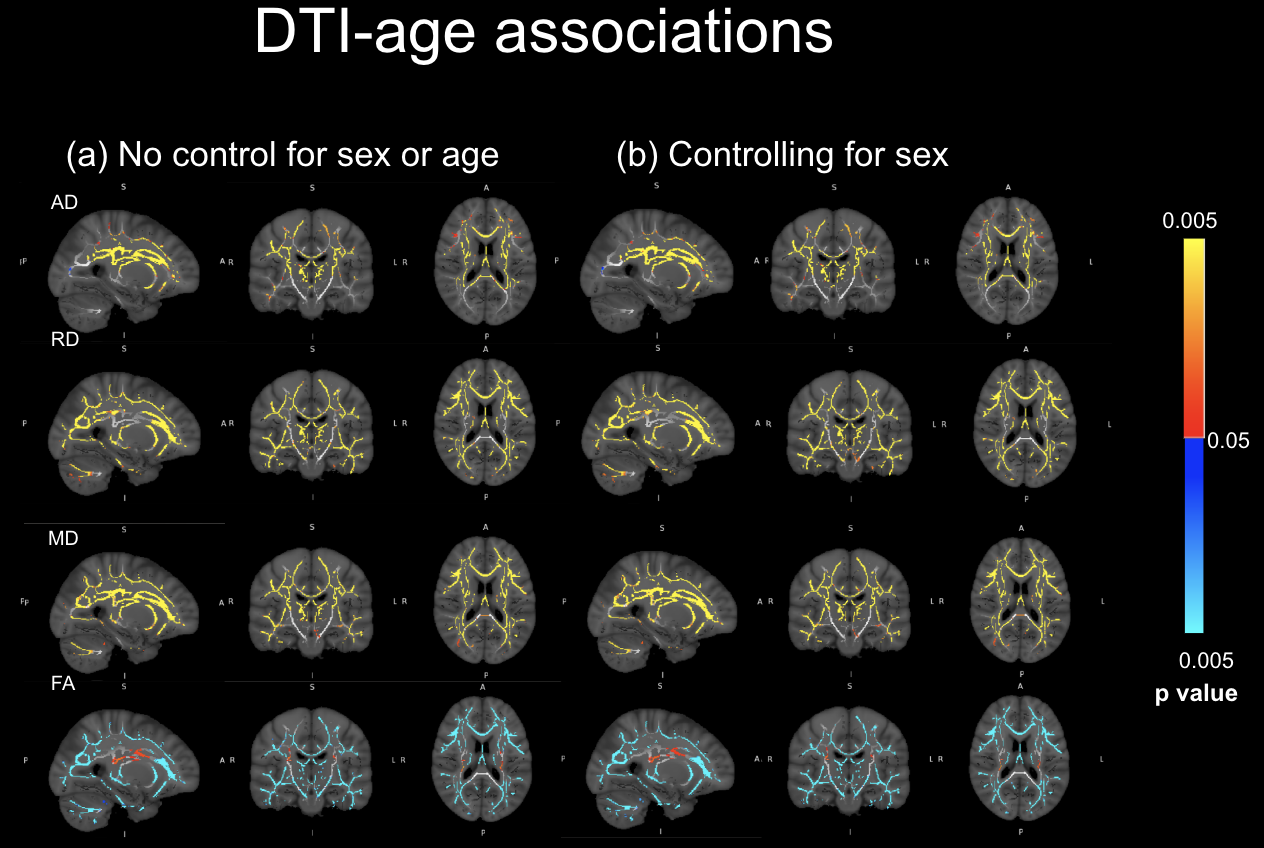

**Figure A1. DTI metrics versus age.** Blue indicates significant negative associations and orange-yellow indicate significant positive associations, corrected for multiple comparisons. The corrected p value of the association is indicated by the colourbar.

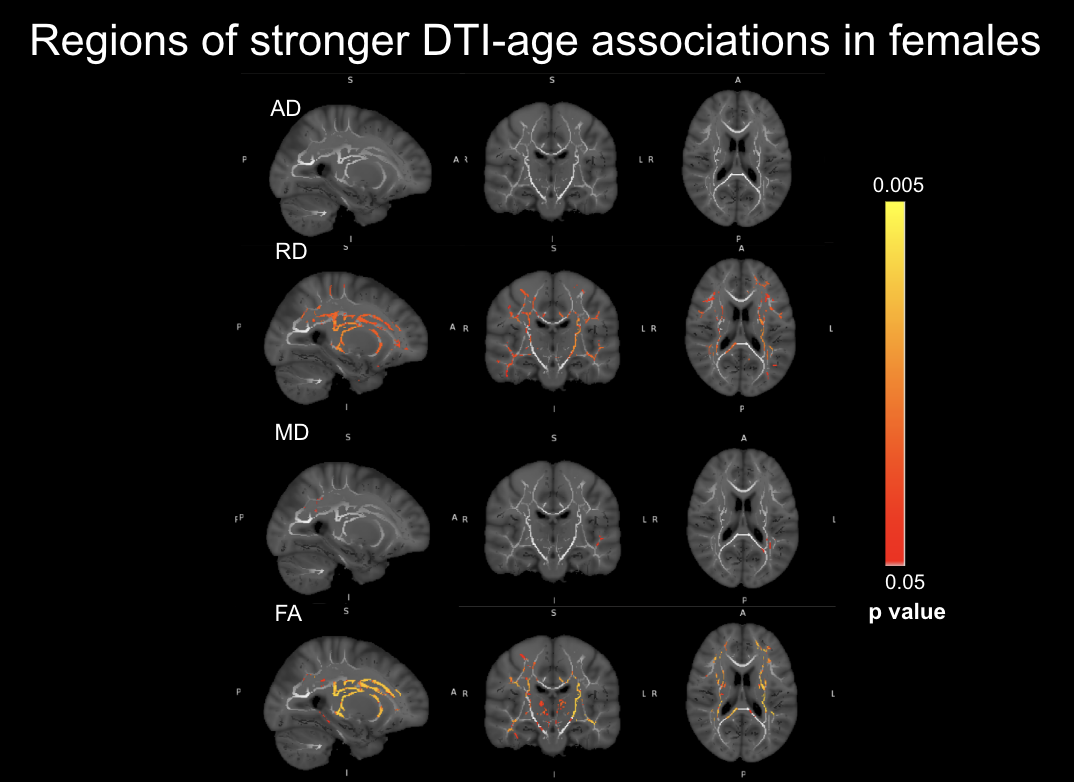

**Figure A2. Regions where DTI-age associations are stronger for females than males.** Displayed are differences in absolute associations. Orange indicates significantly stronger absolute associations in female subjects, corrected for multiple comparisons. The corrected p value of the association is indicated by the colourbar. There are no regions in which the male subjects exhibited a stronger association.

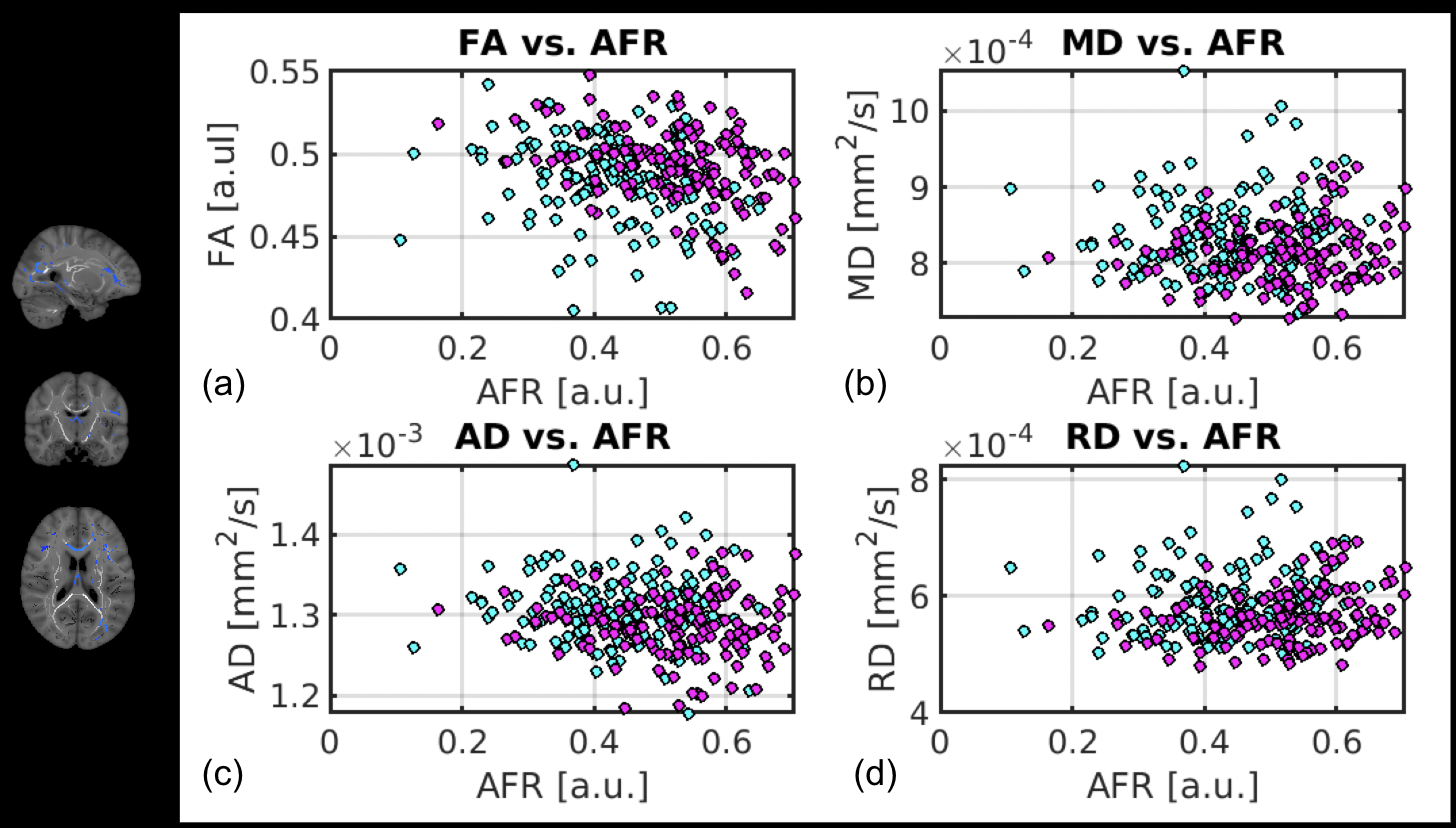

**Figure A3. DTI metrics versus AFR.** All values are taken from the regions of significant association between FA and AFR, controlling for sex (see panel on the left side of figure). AD alone exhibited a significant negative association with AFR in these regions (Fig. 5).

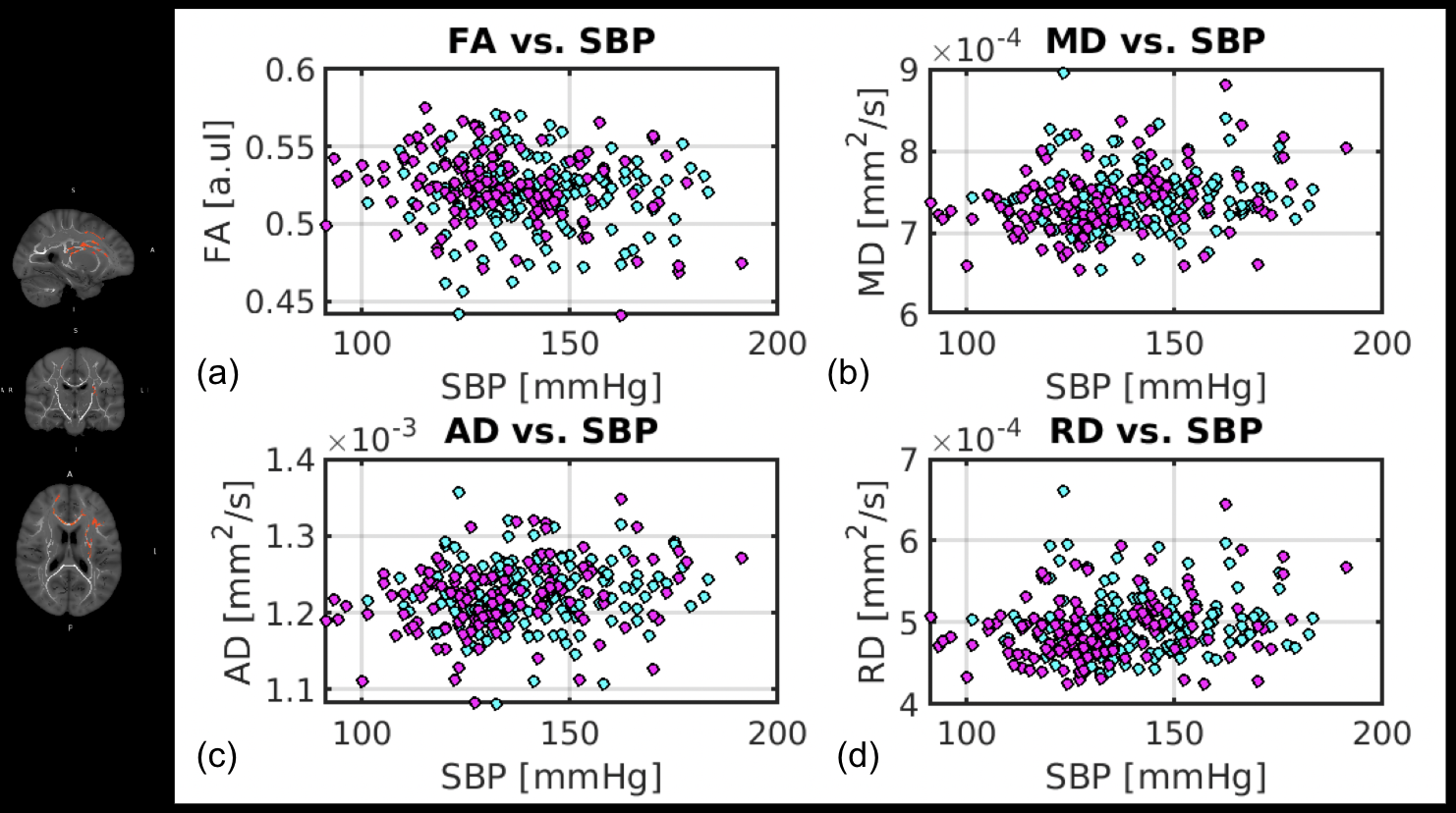

**Figure A4. DTI metrics versus SBP.** All values are taken from the regions of significant association between MD and SBP, controlling for sex (see panel on the left side of figure). MD alone exhibited a significant negative association with MD in these regions (Fig. 6). AD and FA did not exhibit significant association with SBP in these regions.
